# Acclimation, priming and memory in the response of *Arabidopsis thaliana* seedlings to cold stress

**DOI:** 10.1101/848606

**Authors:** Jan Erik Leuendorf, Manuel Frank, Thomas Schmülling

**Affiliations:** Institute of Biology/Applied Genetics, Dahlem Centre of Plant Sciences (DCPS), Freie Universität Berlin, Albrecht-Thaer-Weg 6, D-14195 Berlin, Germany

## Abstract

Because stress experiences are often recurrent plants have developed strategies to remember a first so-called priming stress to eventually respond more effectively to a second triggering stress. Here, we have studied the impact of discontinuous or sustained cold stress (4 °C) on *in vitro* grown *Arabidopsis thaliana* seedlings of different age and their ability to get primed and respond differently to a later triggering stress. Cold treatment of 7-d-old seedlings induced the expression of cold response genes but did not cause a significantly enhanced freezing resistance. The competence to increase the freezing resistance in response to cold was associated with the formation of true leaves. Discontinuous exposure to cold only during the night led to a stepwise modest increase in freezing tolerance provided that the intermittent phase at ambient temperature was less than 32 h. Seedlings exposed to sustained cold treatment developed a higher freezing tolerance which was further increased in response to a triggering stress during three days after the priming treatment had ended indicating cold memory. Interestingly, in all scenarios the primed state was lost as soon as the freezing tolerance had reached the level of naïve plants indicating that an effective memory was associated with an altered physiological state. Known mutants of the cold stress response (*cbfs, erf105*) and heat stress memory (*fgt1*) did not show an altered behaviour indicating that their roles do not extend to memory of cold stress.

---

In temperate or boreal climates low to subzero temperature is an environmental factor influencing plant lifecycle by affecting seed germination, plant growth and the time for flowering. Whereas low temperature stress (< 10 °C) suppresses the growth rate and leaf expansion, freezing temperatures (< 0 °C) lead to a combination of drought stress syndromes, mechanical wounding and finally to plant death^1-3^. Therefore, native plants have developed a strategy, known as cold acclimation^4,5^, to prepare themselves in low temperatures for upcoming freezing temperatures. This process includes the formation and accumulation of cryo-protectants like e.g. soluble sugars^6^, prolines^7,8^, flavonoids or anthocyanin^9,10^, changes in lipid and protein compositions of cellular membranes^11,12^, and major changes in the plant transcriptome and proteome^4,12-14^.

Numerous changes in gene expression accompanying the cold acclimation process are initialized through the C-REPEAT BINDING FACTOR (CBF)-dependent cold pathway. Here, cold-activated CBF transcription factors (CBF1 to CBF3) are inducing cold response genes, which are for example responsible for the biosynthesis of protectants, and *COLD-REGULATED* (*COR*) genes^15^. Well described examples for *COR* genes are *COR15A* and *COR15B*, which are strongly upregulated in a *CBF-*dependent manner and which enhance the freezing resistance by stabilizing the chloroplast membranes when constitutively (over)expressed^16,17^. *CHALCONE SYNTHASE* (*CHS*) is another cold-inducible gene^18^, encoding a key regulatory enzyme of the anthocyanin biosynthetic pathway, but it is not part of the CBF regulon^19-21^. Given the fact that only ∼12 % of the cold-regulated genes are regulated by CBFs^20,21^, one has to assume that also other transcription factors are of importance for plant cold acclimation. For example, recently the transcription factor genes *BZR1*^22^ and *ERF105*^23^ have been shown to be important for regulating freezing tolerance and cold acclimation in *Arabidopsis*. Several plant hormones are also involved in regulating the response to cold, in particular abscisic acid (ABA)^24,25^ and gibberellin (GA)^26^, but also growth hormones such as cytokinin^27^ although its role is debated^28^.

It is important to note that not only long-term cold treatment activates acclimation and an improved response to a second cold stress but also cold treatments as short as 24 h^29^. This modification of a future stress by a previous stress characterizes priming^30,31^. Priming has been described as a resource-saving strategy overcoming the limitations of constitutive cold acclimation^1^. Under conditions of variable temperatures it might be favourable if plants would remember cold stress to be prepared to respond effectively to an eventual future cold stress without investing resources into continued maintenance of the acclimated state. In this way they may use more of the available resources for growth^31^. A variety of mechanisms have been shown to be involved in priming stress responses, ranging from lasting changes in metabolism to modification of transcriptions factors and chromatin changes^31,32^. Priming of the response to cold stress has been shown to be regulated by tAPX^29^ and to be associated with partly accession-specific changes of transcripts and metabolites^14,33,34^. However, genes important for cold memory have as yet not been described while several genes are known that play a role in maintaining a primed state in response to elevated temperatures. These include the transcription factor genes *HEAT SHOCK FACTOR A2* (*HSFA2*)^35^ and *FORGETTER1 (FGT1)*^36^.

The aim of this study was to explore priming and memory of the cold stress response of *Arabidopsis* seedlings in an *in vitro* system. Such a system might be useful as a screening platform to identify cold-priming and -memory mutants. We tested different scenarios with either repetitive or continuous cold treatments of different length on freezing tolerance. Deacclimation and the duration of an eventual priming effect was followed over several days. It was found out that the primability was age-dependent, that repeated nightly priming stimuli had in contrast to continuous cold treatment only a limited capacity of increasing freezing tolerance and that memory to cold was lost after three to four days at the same time when the physiological consequences of priming had disappeared.

## Results

### Age-Dependent Freezing Tolerance in *Arabidopsis* Seedlings

In order to analyse the age dependence of the response of *Arabidopsis thaliana* seedlings to low non-freezing temperatures, we tested first the response of *in vitro* grown seedlings of different ages to a single 16-h night cold treatment. 7-, 14- and 21-d-old *Arabidopsis* seedlings grown under short day conditions were exposed to cold stress (4 °C) during the last night (P1 in Figure 1a). The expression of several cold marker genes (*CBF1, CBF3, COR15A* and *COR78*) all belonging to the CBF regulon, was determined at the end of the night (Figure 1b, c; Supplementary Figure 1). Under control conditions the steady state level of *CBF3* and *COR15A* mRNA was lowest in 7-d-old seedlings compared to 14- and 21-d-old seedlings while it was similar in seedlings of all ages for *CBF1* and *COR78*. During cold treatment the expression of all cold response genes was very strongly induced in seedlings of all ages indicating a similar responsiveness to cold (Figure 1b, c; Supplementary Figure 1).

**Figure 1.**
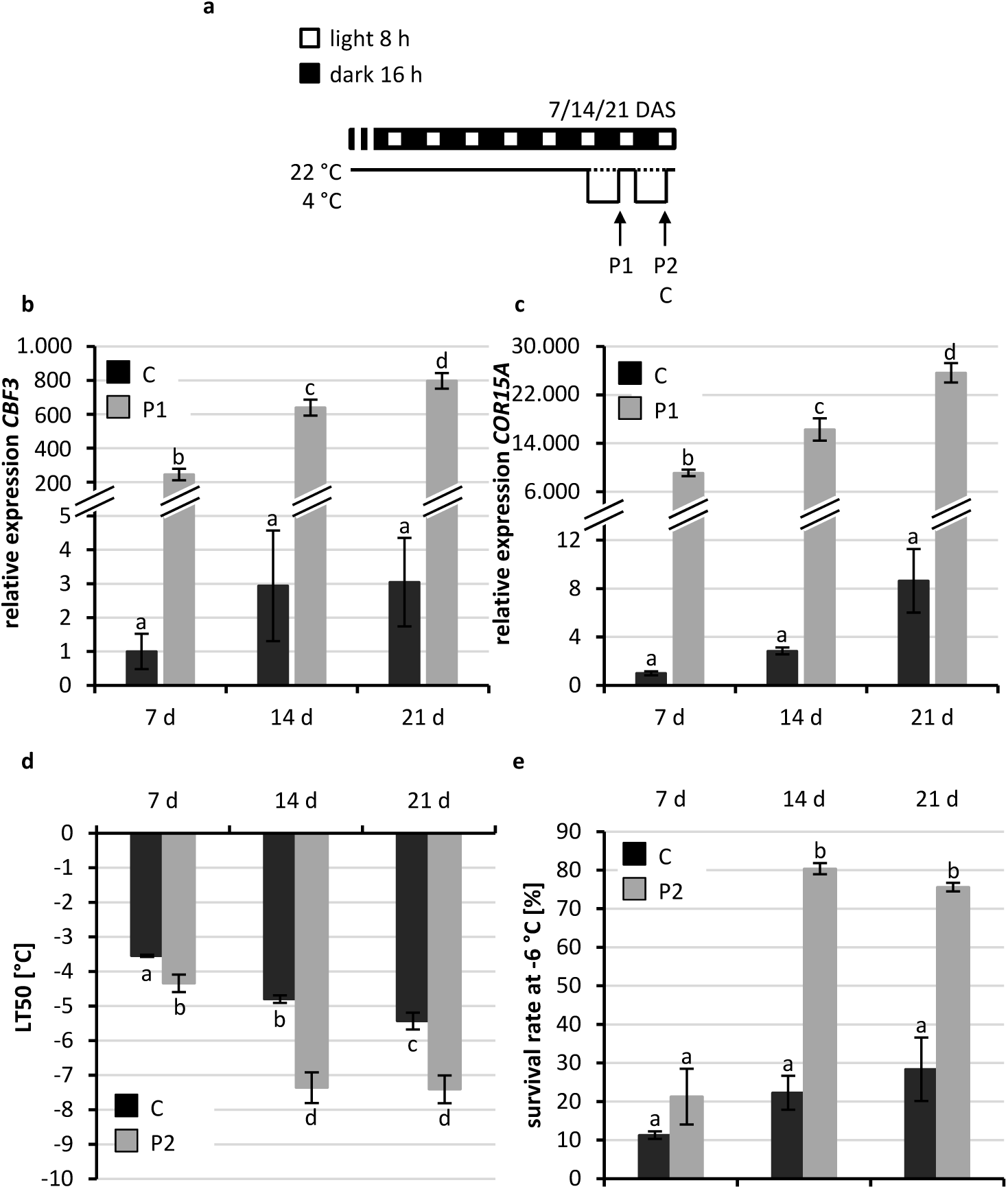
Cold stress response of non-acclimated and cold-primed 7-, 14- and 21-d-old *Arabidopsis* seedlings. (a) Schematic illustration of growth conditions for seedlings tested in b-e. *Arabidopsis* seedlings were grown to the age of 7, 14 or 21 d at 22 °C under short-day conditions (control, C) and cold-primed at 4 °C for the last (P1) or the last two nights (P2). Expression of cold marker genes (b, *CBF3*; c, *COR15A*), (d) electrolyte leakage measurements and (e) survival assay at −6 °C. The freezing tolerance and the cold response of primed plants (P2) was compared to the non-cold acclimated control (C, dotted line in a) of the same age. Significance of differences between conditions was calculated by two-way ANOVA. Identical letters indicate no significant difference (p < 0.05). Error bars in c, e, f represent standard deviation, in d standard error, n ≥ 4. DAS, days after sowing.

Next we compared in cold-treated seedlings of different ages the freezing resistance by determining the temperature causing 50 % damage in electrolyte leakage measurements (LT_50_). Seedlings were exposed to 4 °C during the last two consecutive nights and the LT_50_ value was determined thereafter (P2). Under control conditions, 7-d-old seedlings had a 1.5 - 2 °C lower LT_50_ value as compared to 14- and 21-d-old seedlings indicating their lower resistance to cold stress (Figure 1d). This difference was even larger after two nights of exposure to cold. 7-d-old seedlings showed only a weak response to the cold treatment while 14-d-old and 21-d-old seedlings showed a significantly lowered LT_50_. The older seedlings showed only slight differences in comparison to each other both before and after cold treatment. A survival assay showed a similar result: Older seedlings had a tendency to an increased survival after exposure to −6 °C as compared to 7-d-old seedlings (Figure 1e). Cold treatment prior to the survival assay increased strongly the survival rates of the 14-d-old and 21-d-old seedlings but not in the 7-d-old seedlings (Figure 1e).

Taken together, cold response genes were similarly induced in response to cold in 7-, 14- and 21-d-old seedlings. However, 7-d-old seedlings were not able to develop an increased freezing resistance as did the older seedlings and were thus more vulnerable to freezing stress. We conclude that activation of the CBF/COR pathway is not sufficient for young *Arabidopsis* seedlings to increase their resistance to cold temperatures. Further, *Arabidopsis* seedlings acquire the full competence to respond appropriately to low temperatures only about two weeks after germination.

### Repeated Cold Treatments interrupted by Recovery Phases cause only a limited Acclimation to Cold

Next we asked the question whether it makes a difference if seedlings experience a continuous cold treatment of varying length or repeated treatments with intermittent phases at normal temperature which mimic conditions of cold nights and warmer days occurring during spring. In the latter scenario a first cold stress treatment (the priming stress) would eventually need to be remembered in order to be prepared for the next cold treatment (triggering stress). In all stress regimes the *Arabidopsis* plants were 21-d-old on the day the experiment was ended (Figure 2a).

**Figure 2.**
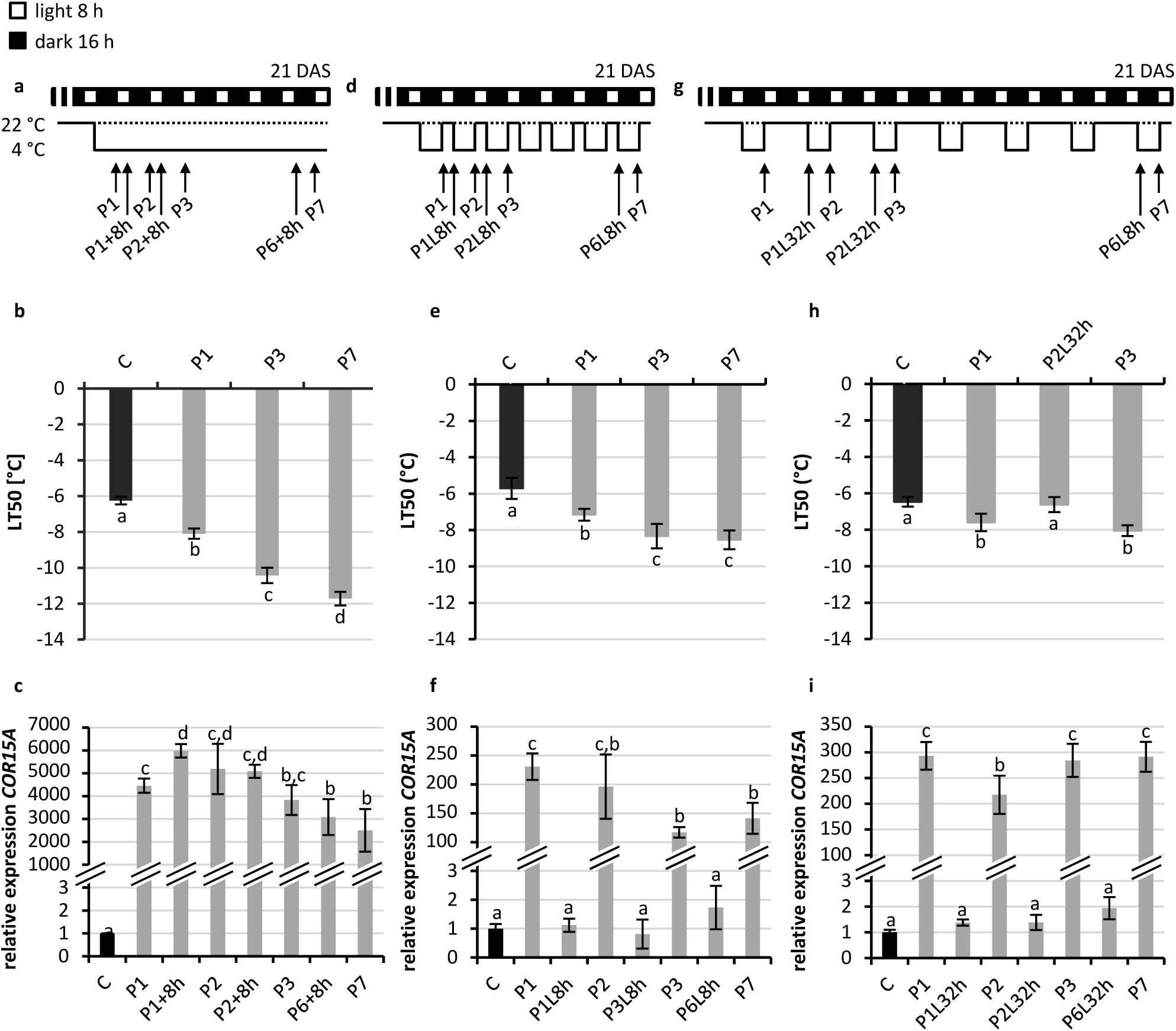
Cold priming response of constantly and repetitively cold-treated *Arabidopsis* seedlings with intermittent recovery phases of various length. (a, d, g) Schematic illustration of the treatments. 21-d-old seedlings treated constantly (a, b, c), every night (d, e, f) or every second night (g, h, i) with 4 °C (b, e, h). Analysis of freezing resistance by electrolyte leakage measurements (LT_50_). (c, f, i) Induction of the cold response gene *COR15a* measured by qRT-PCR. Expression in non-primed control seedlings at the given sampling time (morning or evening) was set to 1. For d) and g) the cold stress response was tested after the given treatments (P1-P7) and after a 8-h or 32-h recovery phase at 22 °C (lag phase; L8h, L32h). For comparison, similar time points were chosen to monitor the response to a constitutive cold stress treatment of seven days (a). Non-cold treated plants (C; dotted line). Significance of differences between the conditions was calculated by one-way ANOVA. Identical letters indicate no significant difference (p < 0.05). Error bars represent standard deviation, n = 4. DAS, days after sowing.

Samples for LT_50_ and gene expression analysis were taken after one day (P1), two days (P2), three days (P3) and seven days (P7) of cold exposure at 4 °C. As expected the constant cold treatment induced a robust acclimation process, whereby the enhancement of the freezing resistance (LT_50_, Figure 2b) strongly correlated with the duration of the treatment reaching at the end an LT_50_ of −11.7 ± 0.38 °C, which is close to the maximum reported for Col-0 grown *in vitro*^37^. For comparison we tested two additional scenarios with up to seven repetitive 4 °C cold treatments during the night (P1 to P7) with intermittent recovery or lag phases at 22 °C for 8 or 32 hours (L8h or L32h; Figure 2d, g) and analysed the freezing tolerance at different time points.

Repetitive cold treatments with a recovery phase of 8 h induced an acclimation response that did not reach by far the level of the constantly cold-treated seedlings (Figure 2e, f). The increase of freezing resistance was significant only during the first stress treatments. The lowest LT_50_ value of −8.3 ± 0.67 °C was reached after three consecutive stress treatments and did not decrease further, possibly because an equilibrium between the priming response and deacclimation was reached (Figure 2e). This LT_50_ value was about 2 °C higher than the - 10.4 ± 0.43 °C reached after three days of continuous cold treatment (Figure 2b). Repetitive cold stress treatments with 32 hours recovery phases led to no further improvement of cold acclimation after the first treatment (Figure 2h). This indicated that the priming effect of the first stress experience that was noted during the first cold treatments with 8 h intermittent recovery phases (Figure 2b) was lost during the longer recovery phase of 32 h. In addition, analysis of the LT_50_ value at the end of the second recovery phase (P2L32h) showed that the freezing resistance at the end of the 32 h-recovery phase was comparable to the basal resistance level of untreated seedlings, i.e. in their naïve state (Figure 2h).

The expression of two cold response marker genes (*COR15A, CHS) w*as measured before and after the second, third and seventh nightly cold stress treatment (Figure 2d and g) as well as at the corresponding time points in the constant cold treatment scenario (Figure 2a). *COR15A* belongs to the CBF regulon while the expression of *CHS* is upregulated by cold independent of the CBF regulon^18,20,21^. In constant 4 °C cold conditions *COR15A* and *CHS* gene expression was highly induced after a single night and decreased only slowly thereafter keeping a high expression level until the end of the experiment after seven days (Figure 2c, Supplementary Figure 2b). Repeated cold treatments induced likewise the cold marker gene expression but the expression was always set back to the basal level of untreated plants (C) after the recovery phases of 8 h or 32 h (Figure 2f and I; Supplementary Figure 2c and e). Interestingly, the induction of the cold marker genes in response to the cold stress treatments with 8 h recovery phase was lower after several cold treatments as compared to the first cold treatment, an effect that was even stronger for *CHS* gene expression than for *COR15A* expression. After three stress treatments the cold marker gene induction reached a stable level which correlated with no further increase in freezing resistance (LT_50_). This effect was not seen in seedlings experiencing a recovery phase of 32 h (Figure 2i). In this scenario cold marker gene expression was induced always to a similar level after the consecutive stress treatments, which correlated also in this case with no further increase in freezing resistance.

These results show that *Arabidopsis* seedlings can store the information about an experienced short cold stress for a short period and remain for at least 8 h but no longer than 32 h in a primed state allowing to establish a higher stress resistance in response to a second triggering cold stress. However, the capacity to build a high level of resistance in response to repeated short (cold) stress treatments was limited, a much stronger effect could only be reached by a continuous cold treatment.

### The Recovery Phase of cold-treated Seedlings is characterized by a fast Decline of Cold Marker Gene Expression and a slower Reduction of the Freezing Tolerance

In order to analyse the duration and the characteristics of the primed state under non-stress conditions we subjected seedlings to a cold-priming treatment (4 °C) of three days and analysed the freezing resistance (LT_50_) and cold marker gene expression immediately after the stress treatment and after two to five days of subsequent recovery (lag) phase at 22 °C (Figure 3a). The priming treatment caused an enhanced freezing resistance and a robust induction of *COR15A, CBF3* and *CHS* (P3, Figure 3b and c; Supplementary Figure 3). After the stress treatment the freezing resistance returned to the level of naïve plants within four to five days (P3L4-5) (Figure 3b). In contrast, the expression level of all three cold response genes dropped to the basal expression level of naïve plants within the first two days after the priming stimulus (P3L2, Figure 3c; Supplementary Figure 3). This indicated that the transcriptional response to cold was lost rapidly after the end of the cold period but that the acquired increased freezing resistance was lost more slowly indicating a cold memory.

**Figure 3.**
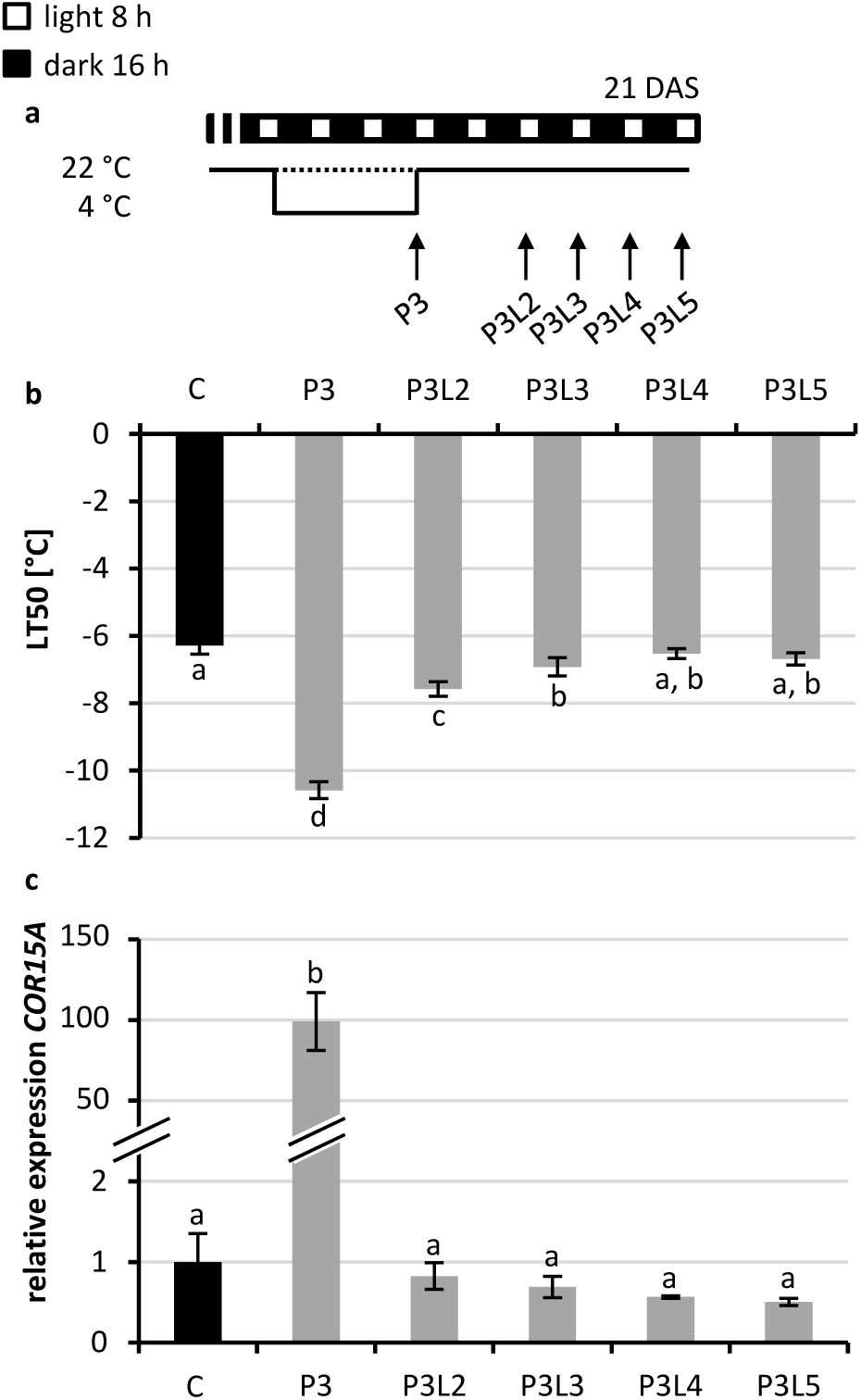
Analysis of the cold memory in *Arabidopsis* seedlings. (a) Schematic illustration of the growth conditions. (b) Analysis of freezing resistance by measurement of electrolyte leakage. (c) Expression of the cold-response gene COR15A (expression under control conditions was set to 1). 21-d-old seedlings were treated for three days with 4 °C cold as a priming stimulus (P3) and subsequently deacclimated for two to five days at 22 °C (P3L2-5). Significance of differences between conditions was calculated by one-way ANOVA (p < 0.05). Identical letters indicate no significant difference. Error bars indicate standard deviation, n = 4. C, non-primed control (dotted line in a); DAS, days after sowing.

### Cold-priming positively affects the Response to a following Cold Stress

Next we tested if the response to a second cold stress, the triggering stimulus (T), could be positively affected by the priming stimulus and if this effect would be dependent or independent of a residual activity of the priming effect. The triggering stimulus was applied for two days after a priming stimulus (three days at 4 °C) and a subsequent lag phase of three to five days (Figure 4a, d, g). The lag phase covered the period during which the consequences of the first cold priming stimulus, an enhanced freezing resistance, was slowly diminished (Figure 3). After a lag phase of three days cold-primed and -triggered plants (P3L3T1, P3L3T2; Figure 4b) showed a significant improvement (−1 °C) of the freezing resistance when compared to only cold-triggered plants (T1, T2) which did not receive the first cold-priming stimulus. After a lag phase of four days only a slight but not significant difference between cold-primed and -triggered and only triggered plants was detected (Figure 4e). At this time point (P3L4T1) results were variable and in some experiments a higher freezing tolerance was noted as compared to triggered only plants (T1). However, the priming effect was completely lost in all experiments after a lag phase of five days (Figure 4h).

**Figure 4.**
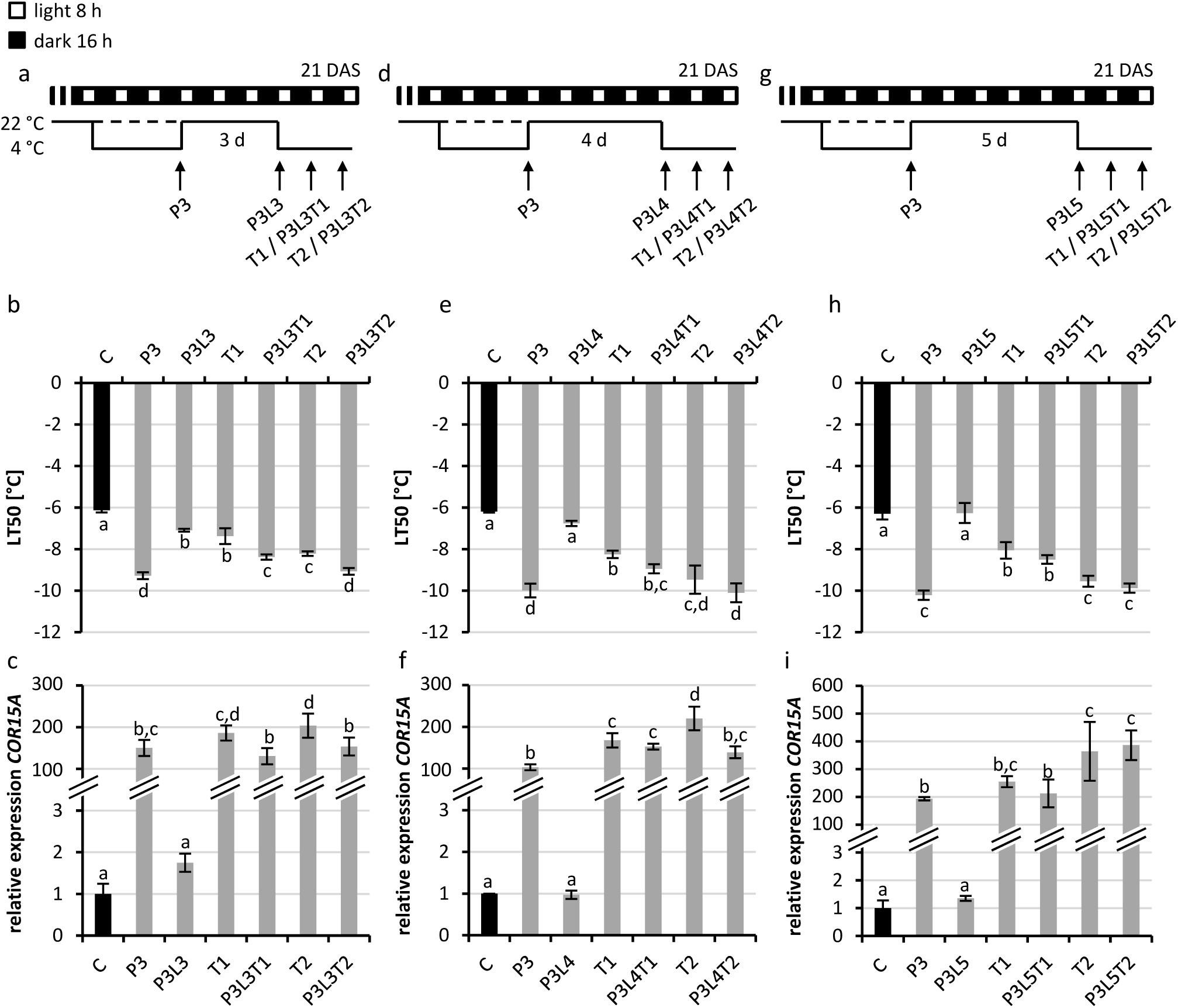
Analysis of the cold priming effect in *in vitro* grown *Arabidopsis* seedlings. (a, d, g) Schematic illustration of the growth conditions. (b, e, h) Analysis of the freezing resistance by electrolyte leakage measurement and analysis of the cold-responsive *COR15A* gene expression (c, f, i) analyzed by qRT-PCR (non-primed control was set to 1). 21-d-old seedlings were analysed after cold-priming at 4 °C for three days (P3), after the following recovery at 22°C (lag phase, L) for three days (a, b, c), four days (d, e, f) or five days (g, h, i) and after an additional 4 °C cold-triggering stimulus for one or two days (T1, T2). Cold-primed and -triggered seedlings (PLT) were compared to only cold-triggered seedlings (T, dashed line in a, d, g). Significance of differences between conditions was tested by one-way ANOVA (p < 0.05). Identical letters indicate no significant difference. Error bars represent standard deviation, n ≥ 3. C, non-primed control; DAS, days after sowing.

Like in the previous experiment (Figure 3) the freezing resistance was still above the basal freezing resistance level of naive plants three (P3L3) but no longer four or five days (P3L4, P3L5) after the end of the cold priming stimulus (Figure 4b, e, h). We conclude that the primed state representing a cold memory leading to an enhanced response to a triggering stimulus was associated with an enhanced freezing resistance during the lag phase. In other words, the priming cold stress experience was forgotten when the freezing resistance reached again the level of naïve plants.

The expression of *COR15A* in this experiment also reflected the previous results. After a strong induction in response to the priming stimulus the expression returned to the basal level of naïve seedlings within the first three days after the lag phase (Figure 4c). In response to the triggering stimulus *COR15A* expression was again highly induced. Interestingly, cold-primed and -triggered plants showed a significantly lower induction of the cold marker gene in response to the cold-triggering stimulus after three days of a lag phase. After a lag phase of five days (P3L5T vs T; Figure 4h) *COR15A* was induced by the triggering stimulus to a similar level in non-primed and cold-primed plants, a pattern that was also seen for the response of *CHS* (Supplementary Figure 4). In sum, the reduced induction of the two cold marker genes in response to the triggering stimulus reflected the primed state of the plants and was in agreement with the improvement of their freezing resistance.

### Known Cold Acclimation and Memory Genes are not required for Cold Memory

It could be that genes involved in cold acclimation are also functionally relevant for maintaining the acclimated state or the deacclimation process, which is linked to memory. Memory genes known from another functional context such as heat stress might have a role in cold memory as well. Therefore, we have next tested the behaviour of a number of mutants in such genes in our experimental setting.

The CBF regulon is the best studied signalling pathway of cold acclimation. We tested whether loss of this pathway negatively influences cold priming and/or memory in *Arabidopsis* seedlings. The *cbfs* triple mutant^38^ (*cbfs*) was used in a cold-priming and - triggering scenario (P3L3T1, Figure 5a) and the freezing resistance was monitored. Untreated and three days cold-primed (P3) wild-type seedlings were analysed as controls (Figure 5b) in parallel. *cbfs* and wild-type seedlings showed a similar freezing tolerance under control conditions but *cbfs* showed significantly less induction of freezing tolerance in response to the cold-priming stimulus, i.e. −9.03 ± 0.32 °C as compared to −10.23 ± 0.32 °C in wild type. This is in agreement with the results of Jia et al. (2016)^38^, who showed that the *cbfs* mutant exhibits a similar basal freezing tolerance but less acclimation capacity in response to cold as compared to wild type. Additionally we observed in *cbfs* a less strong induction of the *COR15A* gene in response to the cold-priming stimulus (Supplementary Figure 5) which is again in agreement with the published data ^38^.

**Figure 5.**
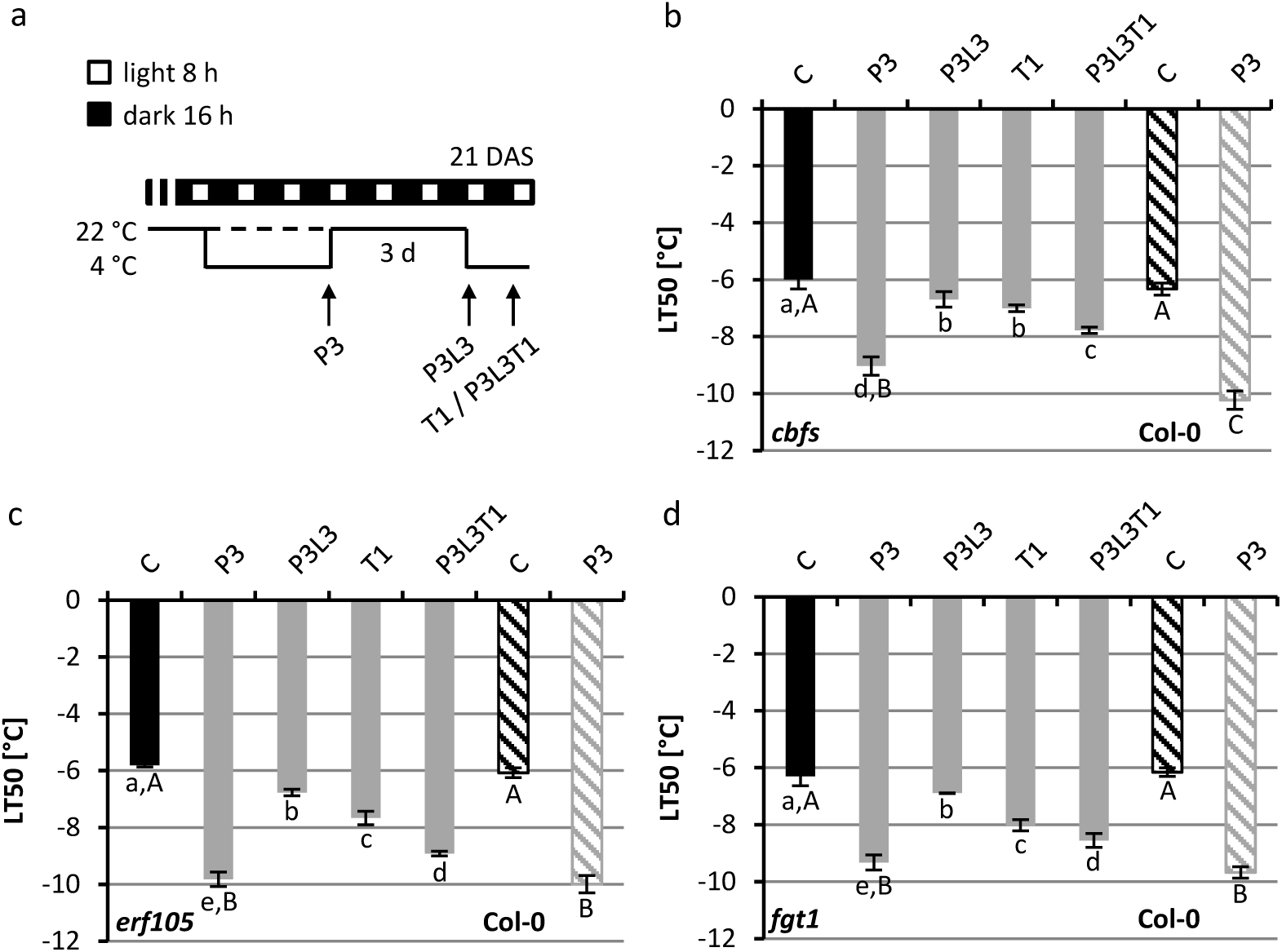
Cold priming and memory in cold response and memory mutants. (a) Schematic illustration of the growth conditions. Analysis of freezing resistance by measurement of electrolyte leakage (LT_50_) in the *cbfs* (b), *erf105* (c) and the *fgt1* mutant (d). 21-d-old seedlings were analysed after three days cold-priming at 4 °C (P3), after a subsequent lag phase of three days at 22 °C (L3) and after an additional 4 °C cold triggering stimulus for one day (T1). Cold-primed and -triggered seedlings (P3L3T1) were compared to only cold-triggered plants (T1, dashed line). Results for in parallel tested wild-type Col-0 control plants are shown as striped columns. Significance differences of the mutant genotype between conditions were tested by one-way ANOVA (small letters), between the mutant genotype and Col-0 (C and P3) by two way ANOVA (capital letters). Identical letters indicate no significant difference (p < 0.05). n = 4. C, control; DAS, days after sowing.

After three days of recovery *cbfs* mutant plants showed a residual freezing resistance level slightly but significantly above the control at the end of the three day period (P3L3) similar to wild type (Figure 4b). Like in the wild type (Figure 4b), a following triggering stimulus caused in *cbfs* a significantly higher induction of freezing resistance in comparison to only cold-triggered plants (Figure 5b). Thus the *cbfs* mutant behaved generally similar like wild type in this priming and triggering scenario, although with a weaker response to cold. It can be concluded that the three *CBF* genes have no essential role in maintaining the priming-induced cold memory.

Another transcription factor gene relevant for freezing tolerance and acclimation is *ERF105*^23^. The behaviour of *erf105* in response to our priming and memory scenario (P3L3T1) was similar as the response of the wild type when comparing cold-primed and triggered or only cold-triggered plants (Figure 4b and Figure 5c). This indicated that ERF105 has no strong role in younger plants and that mutation of ERF105 does apparently not alter the memory to a cold priming event.

Seedlings with a reduced cytokinin signalling have been reported to display an altered response to cold^27^. However, in the tested scenario (P3L5T1) the double cytokinin receptor mutant *ahk2 ahk3* did not differ from wild type (Figure 4h and Supplementary Figure 6) showing that a reduction of cytokinin signalling did neither alter the basic freezing resistance nor the primability of the tested plants.

Finally we tested the *forgetter1 (fgt1)* mutant, which has been described as a memory mutant defective in keeping active the primed state established in response to a heat stress stimulus^36^. In the standard cold priming and memory scenario (P3L3T1) *fgt1* responded like wild type: both genotypes showed a similar cold priming response, residual enhanced freezing tolerance after three days lag phase and enhanced freezing tolerance in cold-primed and -triggered plants compared to only cold-triggered plants (Figure 5d). These results do not support an important role for FGT1 in the memory to cold stress.

## DISCUSSION

At first, the results show that the cold response and cold acclimation of *Arabidopsis* were dependent on the plant age. Both, electrolyte leakage experiments and survival assays indicated that *in vitro* grown 7-d-old seedlings were not or less able to enhance their freezing resistance after two days at 4 °C (Figure 1), which is consistent with a recent report^39^. Because 7-d-old seedlings grown under short day conditions have essentially still only two cotyledons and 14- to 21-d-old seedlings have developed true leaves it is likely that the competence to develop an enhanced freezing resistance in response to cold treatment is coupled to the development of true leaves. Interestingly, despite their inability to sufficiently enhance their freezing resistance the expression of *CBF3* and *COR15A* genes was also strongly induced in response to cold in 7-d-old seedlings. This demonstrated that these seedlings are able to sense cold temperatures and activate the CBF pathway. However, activation of this pathway appears not to be sufficient for cold acclimation in 7-d-old seedlings but other pathways are required which might not yet be established at this developmental stage. Also cotyledons of *in vitro* grown spinach seedlings showed a tendency of a reduced freezing resistance and less pronounced cold acclimation capacity when compared to true leaves^40^. Wanner and Junttila^41^ proposed that the reduced freezing resistance of *Arabidopsis* cotyledons and their incapability to acclimate to cold stress may be linked to decreased production of photosynthetic assimilates acting as cryo-protectants. However, this alone could not explain the difference in freezing resistance because overproduction of soluble sugars in transgenic tobacco plants did not correlate with an enhanced freezing resistance^42^. Instead, other more complex metabolic and transcriptomic changes associated with an enhanced freezing tolerance might be relevant^14^.

Comparison of the acclimation process in constantly and repetitively 4 °C cold-treated *Arabidopsis* seedlings revealed that *Arabidopsis* is able to acclimate to low temperatures in response to both treatments (Figure 2). In case of repetitive cold treatments, a decisive factor for the degree of acclimation was the length of the intermittent lag phase between treatments. An 8-h-long lag phase was short enough for continuous support of the acclimation process resulting in a stepwise enhancement of the freezing resistance (LT_50_) over three repetitive cold treatments. In contrast, three subsequent cold treatments with an intermittent lag phase of 32 h resulted in a freezing resistance comparable to one achieved by a single treatment. We conclude that information about a nightly cold experience is stored for at least 8 h but not for 32 h. Changes induced by a first treatment may be used as a basis to continue the acclimation process when another phase of cold stress occurs. However, after the changes causing a lowered LT_50_ had disappeared, the memory of stored information was apparently lost and the plants responded to subsequent cold stress treatments like naïve plants receiving the first cold stress. Noteworthy, a constant cold treatment was significantly more effective than repetitive nightly treatments in increasing freezing tolerance which could be due to the rapid beginning of deacclimation during intermittent lag phases not allowing LT_50_ to reach beyond a certain level. The eventual role of the time of day and the circadian clock in regulating the response to short exposures to cold should be considered in future experiments^43,44^.

Interestingly, the response to cold of the tested marker genes *COR15A* and *CHS* was reduced after the freezing resistance was improved by repetitive or constant cold stress treatments (Figure 2f, Supplementary Figure 2d), which is consistent with differential behaviour of cold response genes after repeated cold treatment^29,39^. In repetitive drought stress treatments genes were classified as memory genes if their response to a repeated stress treatment caused a higher or lower change of transcript abundance as compared to the first stress treatment^45,46^. The increase of transcript abundance was explained by the stress-induced establishment of H3K4me3 histone modification which stays high after the previous stress declined and positively affects transcription in the subsequent stress condition^45^. In our stress scenario *COR15A* and *CHS* behaved like memory genes but showing a decreased response to recurrent cold treatment. Interestingly, *COR15A* was shown to be a target of H3K27me3 histone modification, a histone mark negatively influencing gene expression^47,48^. It remains to be tested whether altered gene expression in response to the repeated stress regime used here is associated with histone modification or whether mechanisms modulate the effect of repeated stress on gene expression.

The next question we addressed was how long after a cold treatment plants would keep the enhanced freezing resistance and for how long an additional cold treatment would cause an enhanced response indicating that these plants were still primed and remembered the past treatment. Under our experimental conditions the LT_50_ value of naïve plants was reached three to four days after the end of a 3-d-long cold treatment while the expression of *COR15A* was returned to the initial naïve level already much earlier (Figure 3). This timing of deacclimation was similar as reported for other cold-priming and deacclimation scenarios^33,47^.

Interestingly, when primed plants were challenged with a second triggering cold treatment only those plants responded with an enhanced LT_50_ that still showed measurable physiological consequences of the first cold treatment, i.e. a lower LT_50_ than naïve plants (compare P3L3T1 vs P3L3 and T1 and P3L5T1 vs P3L5 and T1 in Figure 4). After the freezing resistance of cold-stressed plants reverted to the basal unstressed state, no enhancement of the freezing resistance in response to the second cold-triggering stimulus was found when compared to unprimed, only cold-triggered plants. This indicated that memory of priming lasted for about three days and was associated with an altered physiological state of the primed plants (Figure 6). The duration of cold memory found here was a bit shorter than a cold memory of five to seven days reported for soil-grown plants, which however were older and/or primed for a longer period^14,39^.

**Figure 6.**
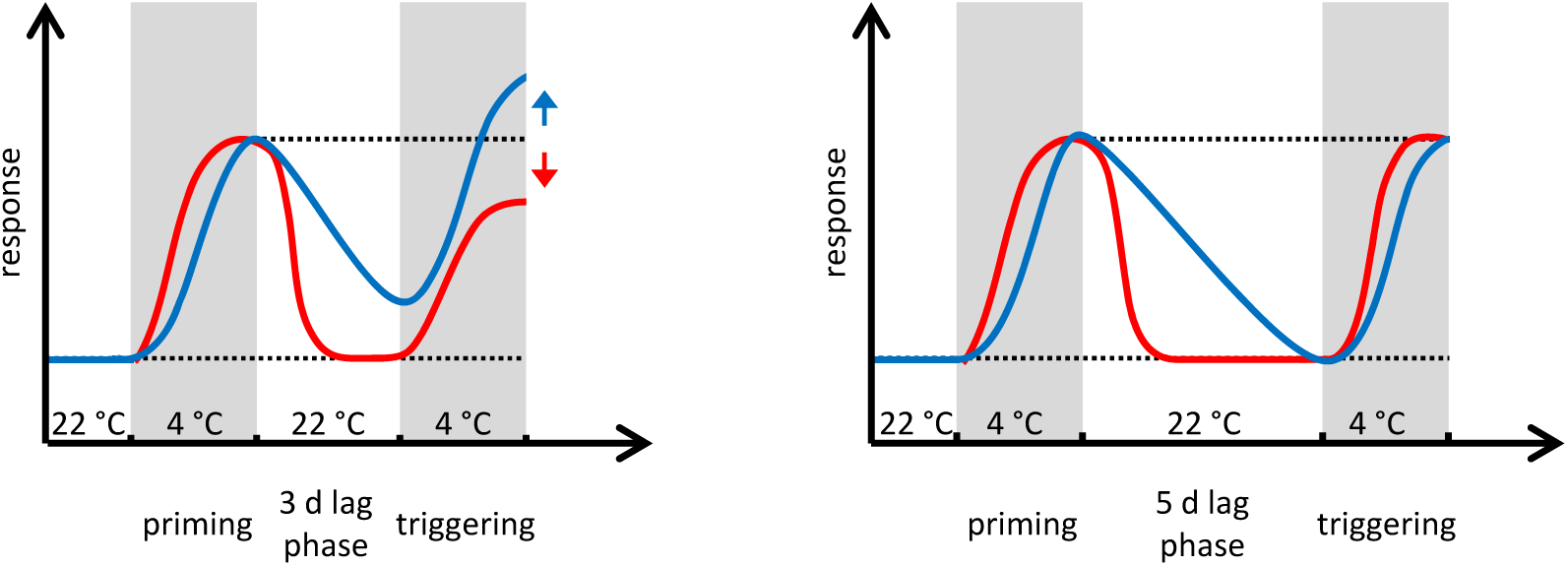
Dependence of cold memory on the cold-primed phenotype. An enhanced freezing resistance (blue line) and a reduced induction of cold responsive genes (e.g. *COR15a*; red line) in response to a cold-triggering stimulus indicate cold memory in primed plants (left). Right: After the freezing resistance had reached the original level the priming effect disappeared and a triggering stimulus induces a cold response similar as in naïve plants. Dotted black lines indicate the unstressed state or level of enhanced priming response.

We have tested in our system the behaviour of several mutants of genes known to be involved in cold acclimation including *cbfs*^38^, *erf105*^23^ and *ahk2 ahk3*^27^. *Cbfs* showed a slightly weaker response to priming while *erf105* and *ahk2 ahk3* responded similar to wild type. This indicated a limited function of CBFs in response to short cold stress in seedlings and suggested that ERF105 is required only later in development^23^. This result supports the notion that the CBF-dependent signalling pathway is not exclusively responsible for the cold stress response in *Arabidopsis* but that additional pathways must exist^20,21,23,38,49^. Neither these mutants nor the heat shock memory gene mutant *fgt1* showed an altered duration of the cold memory under our experimental conditions. We conclude therefore that there must be other factors that are important to respond to low temperature and establish and/or maintain a memory to cold stress. We propose that the protocol established here will be helpful to systematically test additional candidate genes of cold stress memory and/or establish genetic screens that will lead to the discovery of factors involved in cold stress memory. For example, such screens may involve the search for altered expression characteristics of cold response genes or survivors of temporal harsh cold treatments, similar to the successful approaches taken to study the memory of heat stress^36,50^.

## Methods

### Plant Material

*Arabidopsis thaliana* accession Col-0 was used as wild type. Seeds of the *forgetter1-1* (*fgt1-1*)^36^ and the *cfb* triple mutant (*cfbs*)^38^ were kindly provided by Isabel Bäurle and Shuhua Yang. The *ahk2 ahk3*^51^ and *erf105*^23^ mutants were published previously.

### Growth Conditions and Cold Treatment

Plants were grown *in vitro* on half-strength MS Media (w/o vitamins and w/o sugar) with 0,8 % agar under short day (SD) conditions (8 h light/16 h dark) without a stratification period in a climate chamber (Percival) at 22 °C with 120 µmol m^-2^ s^-1^ white light. The 4 °C cold priming stimuli were applied in an identical climate chamber either during the 16-h night or constantly during day and night at the indicated seedling age. During cold treatment the light intensity was lowered to 65 µmol m^-2^ s^-1^ to avoid damage of the photosynthetic apparatus. If not mentioned otherwise samples for electrolyte leakage measurements and expression analysis were taken immediately before the beginning of the night or one hour after the end of the night.

### RNA Extraction and qRT-PCR Analysis

Total RNA was extracted from total seedlings (see growth conditions) using the NucleoSpin RNA Plant Kit (Macherey & Nagel, Düren, Germany) according to the manufacturer’s instructions. Additionally DNase digestion was applied (Thermo Scientific™) to prevent contamination with gDNA. For qRT-PCR analysis, cDNA was synthesized with SuperScript III reverse transcriptase (Invitrogen, Thermo Scientific™) using 1 µg of total RNA. qRT-PCR was carried out as described in Cortleven *et al*. (2016) using gene specific primers (Supplementary Table1) and the CFX96 touch real time PCR detection system (Bio Rad). Gene expression data were normalized against three different reference genes (*ACTIN2*, At3g18780; *Serine/threonine-protein phosphatase 2A regulatory subunit*, At3g25800, and *Transcription initiation factor subunit TFIID*, AT4g31720).

### Survival Assay

For the survival assay surface sterilized seeds were germinated on half-strength MS medium (w/o vitamins and w/o sugar) in a standard petri dish and cultivated for the given time under short day conditions. Seedlings on MS media were tempered for one hour at −1.5 °C in a cultivation chamber (Binder KB 53, Tuttlingen, Germany). Ice crystals were applied to start the crystallization process. The seedlings were cooled down at a cooling rate of −1.5 °C h^-1^ to the final temperature at −6 °C and exposed for two hours. Afterwards, plates were incubated over night at 4 °C, excessive water was discarded and plates were then transferred to standard short day conditions. Two days later, survival was scored by determining visually the condition of the shoot apical meristem and the two youngest leaves. For each tested condition four replicates, with 27-30 seedlings each, were analysed.

### Electrolyte Leakage

Electrolyte leakage as a measure of freezing tolerance was determined for *in vitro*-grown seedlings (root and shoot tissue) over a temperature range from −2 to −12 °C for non-acclimated plants or for plants monitored during the lag phase and from −4 to −14 °C for cold-acclimated plants using a cooling rate of −4 °C h^-1^ as described by^52^. Four replicates were analysed, each of the replicates consisted of six temperature points with a pool of 6-8 seedlings harvested from a separate petri dish (∼50 seedlings). For every replicate a sigmoidal curve was fitted to the leakage values of the six temperature points and the temperature of 50 % electrolyte leakage (LT_50_) was determined using Microsoft Excel and its add-in solver function. Results showing the mean of the LT50 value of the four replicates and their standard deviation.

### Statistical Analyses

Statistical significant differences were calculated by One- or Two-Way ANOVA, followed by Tukey’s Post Hoc test for multiple comparisons. Analysis was carried out by using the software GraphPad Prism7 (GraphPad Software, Inc., La Jolla, USA).

## Supporting information

Manuscript file

## Data Availability

All data generated or analysed during this study are included in this published article and its Supplementary Information files.

## ACKNOWLEDGEMENTS

We thank Ariane Hohenstein and Venja M. Röber for skillful technical assistance. We acknowledge funding by Deutsche Forschungsgemeinschaft (Collaborative Research Centre 973, www.sfb.973).

## Author Contributions

J.E.L and T.S. conceived the project and J.E.L. performed the experiments with contributions from M.F. J.E.L and T.S. wrote the paper, all authors reviewed the manuscript.

## Additional Information

### Supplementary information

accompanies this paper

