## Supplementary material for "Acclimation, priming and memory in the response of *Arabidopsis thaliana* seedlings to cold stress": Manuscript file

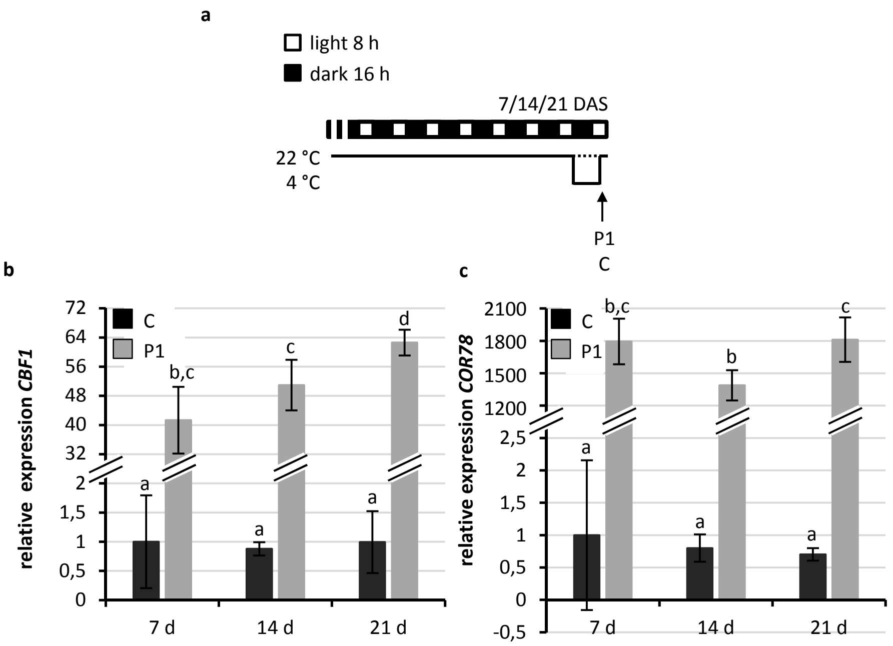


**Supplementary Figure 1.** Cold stress response of non-acclimated and cold-primed 7-, 14- and 21-d-old *Arabidopsis* seedlings. (a) Schematic illustration of growth conditions for seedlings tested in b and c. *Arabidopsis* seedlings were grown to the age of 7, 14 or 21 d at 22 °C under short-day conditions (control, C) and cold-primed at 4 °C for one night (P1). The expression of cold responsive genes (b, *CBF1*; c, *COR78*) in response to cold (P1) was compared to the non-cold acclimated control (C, dotted line in a) of the same age. Significance of differences between conditions was calculated by two-way ANOVA. Identical letters indicate no significant difference ($p < 0.05$). Error bars in represent standard deviation, $n \geq 4$. DAS, days after sowing. This figure is complementary to Figure 1.

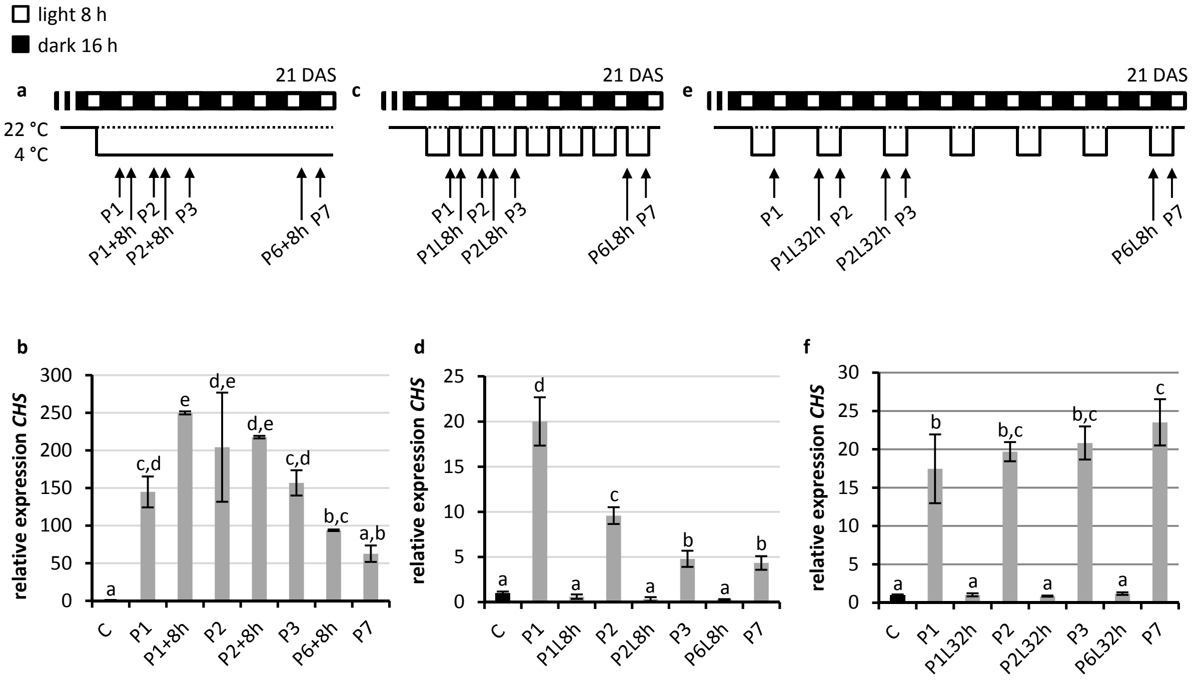


**Supplementary Figure 2.** Cold priming response of constantly and repetitively cold-treated *Arabidopsis* seedlings with intermittent recovery phases of various length. (a, d, g) Schematic illustration of the growth conditions. 21-d-old seedlings treated constantly (a, b, c), every night (d, e, f) or every second night (g, h, i) with 4 °C (b, e, h). Expression of the *CHS* gene analysed by qRT-PCR (c, f, I; C was set to 1). For (c) and (e) the cold stress response was tested after the given treatments (P1-P7) and after a 8-h or 32-h recovery phase at 22 °C (lag phase; L8h, L32h). For comparison, similar time points were chosen to monitor the response to a constitutive cold stress treatment of seven days (a). Non-primed plants (C; dotted line). Significance of differences between the conditions was calculated by one-way ANOVA. Identical letters indicate no significant difference ($p < 0.05$). Error bars represent standard deviation, n = 4. DAS, days after sowing. This figure is complementary to Figure 2.

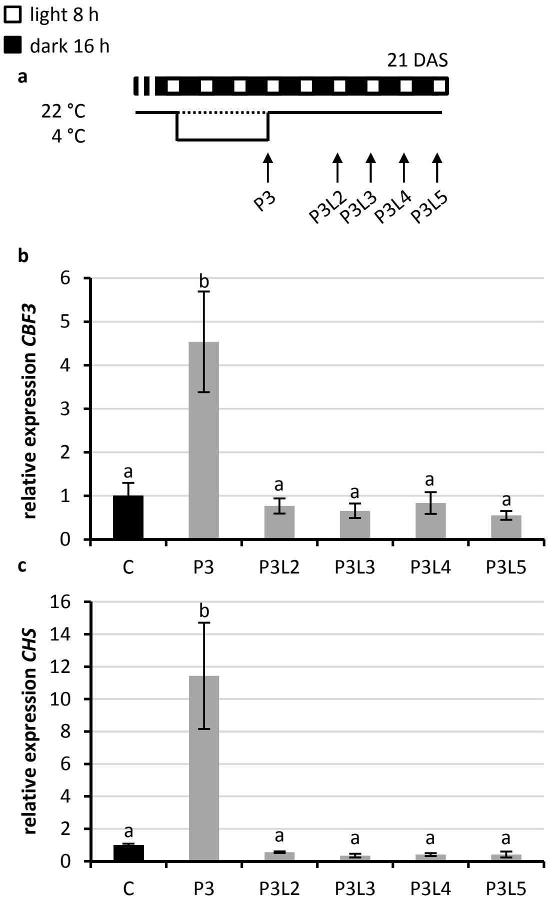


**Supplementary Figure 3.** Analysis of the cold memory in *Arabidopsis* seedlings. (a) Schematic illustration of the growth conditions. Expression of the cold-response gene *CBF3* (b) and *CHS* (c). The expression under control conditions was set to 1. 21-d-old seedlings were treated for three days with 4 °C cold as a priming stimulus (P3) and subsequently deacclimated for two to five days at 22 °C (P3L2-5). Significance of differences between conditions was calculated by one-way ANOVA ($p < 0.05$). Identical letters indicate no significant difference. Error bars indicate standard deviation, n = 4. C, non-primed control (dotted line in a); DAS, days after sowing. This figure is complementary to Figure 3.

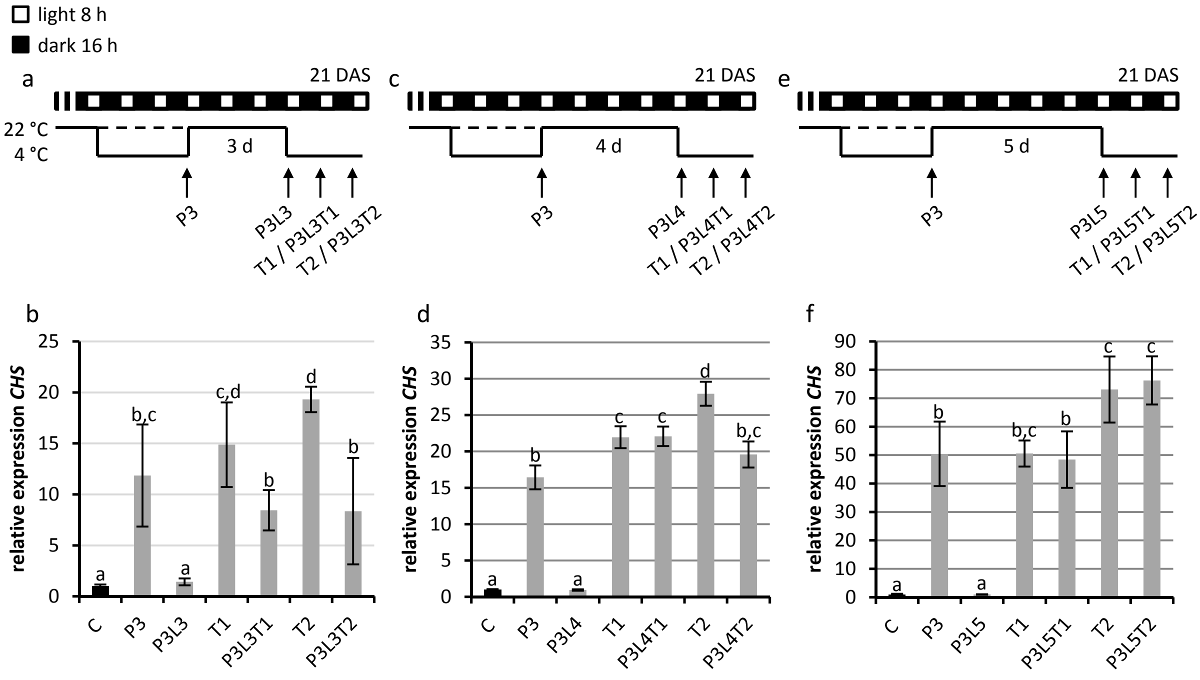


**Supplementary Figure 4.** Analysis of the cold priming effect in *in vitro* grown *Arabidopsis* seedlings. (a, c, e) Schematic illustration of the growth conditions . (b, d, f) Expression of the cold-responsive *CHS* gene analyzed by qRT-PCR (C was set to 1). 21-d-old seedlings were analysed after cold-priming at 4 °C for three days (P3), after the following lag phase at 22 °C (L) for three days (a, b), four days (c, d) or five days (e, f) and after an additional 4 °C cold triggering stimulus for one or two days (T1, T2). Cold-primed and -triggered seedlings (PLT) were compared to only cold-triggered seedlings (T, dashed line in a, c, e). Significance of differences between conditions was tested by one-way ANOVA ($p < 0.05$). Identical letters indicate no significant difference. Error bars represent standard deviation, $n \geq 3$. C, non-primed control; DAS, days after sowing. This figure is complementary to Figure 4.

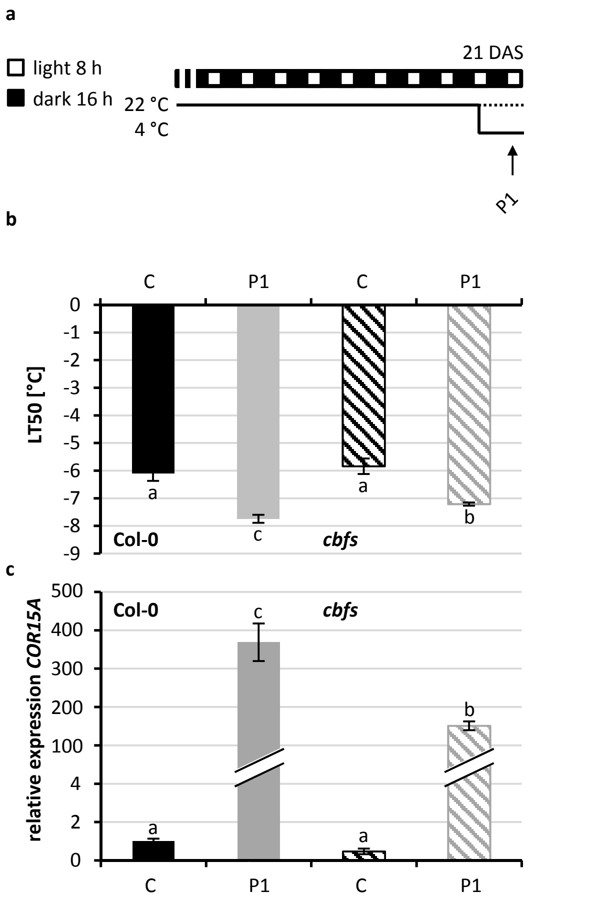


**Supplementary Figure 5.** Cold response of the *Arabidopsis* cold pathway mutant *cbfs*. (a) Schematic illustration of the growth conditions. (b) Analysis of the freezing resistance ($LT_{50}$) of one day cold-primed (P1) or non-cold primed 21-d-old seedlings. (c) Induction of the cold response gene *COR15A*. Significance of differences between conditions or genotypes was tested by two-way ANOVA, n = 4. Error bars indicate standard deviation. Non-cold primed control (C, dotted line in a); DAS, days after sowing.

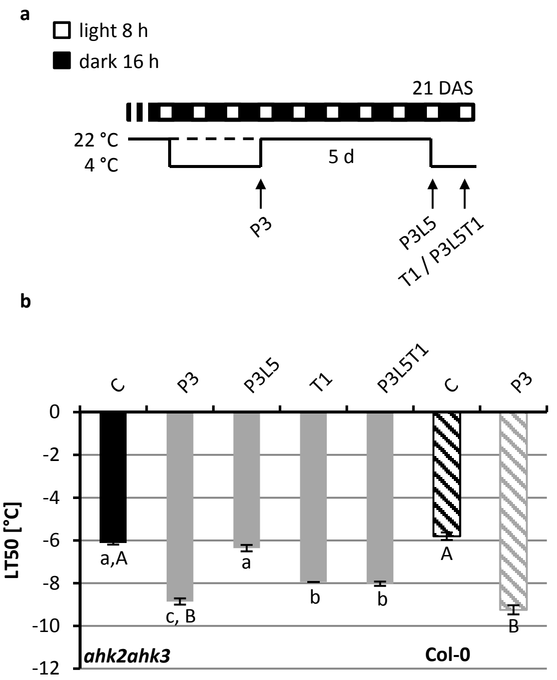


**Supplementary Figure 6.** Analysis of cold priming and memory in the cytokinin receptor mutant *ahk2 ahk3.* Schematic illustration of the growth conditions. (b) Analysis of freezing resistance by measurement of electrolyte leakage ($LT_{50}$). 21-d-old seedlings were analysed after three days cold-priming at 4 °C (P3), after a subsequent lag phase of five days at 22 °C (L5) and after an additional 4 °C cold triggering stimulus for one day (T1). Cold-primed and -triggered seedlings (PLT) were compared to only cold-triggered plants (T, dashed line). Results for in parallel tested wild-type Col-0 control plants are shown as striped columns. Significance differences of the *ahk2 ahk3* mutant genotype between conditions were tested by one-way ANOVA (small letters), between the mutant genotype and Col-0 (C and P3) by two way ANOVA (capital letters). Identical letters indicate no significant difference ($p < 0.05$). n = 4. C, control; DAS, days after sowing.

**Supplementary Table 1.** Oligonucleotides used for qRT-PCR analysis.

| Gene | Gene ID | Forward primer | Reverse primer |
|---|---|---|---|
| *CBF3/DREB1A* | AT4G25480 | CTGATCAATGAACTCATTTTCTGC | TTAATAACTCCATAACGATACG |
| *COR15A* | AT2G42540 | AACGAGGCCACAAAGAAAGC | CAGCTTCTTTACCCAATGTATCTGC |
| *CHS/TT4* | AT5G13930 | TATCGCTAAGGATCTCGCC | GGTGGGCTATCCAGAAGAG |
| *ACT2* | At3g18780 | CTTGCACCAAGCAGCATGAA | CCGATCCAGACACTGTACTTCCTT |
| *TFII* | AT4G31720 | GAATCACGGCCAACAATC | ACTCTTAGCCAAGTAGTGCTCC |
| *PP2A* | At3g25800 | CGTGCGGTGTCTCTTCTT | TTTGATGTTTGGAACTCTGTCTTT |
